# Neural correlates of individual differences in aversion to risk and choice inconsistency

**DOI:** 10.1101/2021.05.10.443362

**Authors:** Manon E. Jaquerod, Alessandra Lintas, Gabriele Gratton, Kathy A. Low, Philippe N. Tobler, Alessandro E. P. Villa

## Abstract

When making financial choices, most people prefer smaller but more certain gains to larger but more uncertain ones with the same expected value (risk aversion). However, attitudes toward risk may vary greatly also within individuals (choice inconsistency). To examine the brain dynamics implementing risky and inconsistent decisions, we recorded event-related brain potentials (ERPs) from 24 adults engaged in a task requiring choices between certain but smaller gains and uncertain but larger ones. Choice consistency and risk aversion were quantified for each individual. Participants were classified into three groups according to their attitude toward risk. Relative neutrality to risk was accompanied by a higher consistency across trials than risk aversion or proneness. Choice consistency was related to the amplitude of the positive ERP peaking near 200 ms after stimulus onset (P200), while risk aversion was related to modulation of the medial frontal negativity (MFN) and to the amplitude of a late positive potential (LPP). Late ERP activity was related to the modulation of value signals by risk levels and associated with individual differences in behavior. Overall, this study suggests that individual differences in attitude toward risk and choice consistency are associated with distinct brain dynamics.

## 1 Introduction

Individuals typically exhibit aversion to risks when possible financial outcomes are gains but seek risks when outcomes are potential losses (Kahneman and Tversky, 1979). For instance, many individuals prefer a smaller certain gain over a larger but uncertain one with the same expected value. However, on top of these overall tendencies, large individual differences in behavior can be observed (Hey, 2001; Frey et al., 2018). These variations in financial choices are thought to be the outcome of a subjective valuation process, where the value of risky outcomes differs across individuals (Christopoulos et al., 2009). Moreover, most decision-making theories assume that preferences rely on *stable* valuations of each alternative choice. However, in reality, subjective valuation is a *dynamic* process (Louie et al., 2013). The concept of choice inconsistency (i.e., a choice that is not consistent with a person’s declared preferences) was introduced to capture the dynamic nature of the valuation process in repeated decisions (Grether and Plott, 1979; Kachelmeier and Shehata, 1992; Seidl, 2002; Tsetsos et al., 2010; Harless and Camerer, 1994; Hey and Orme, 1994) leading to the development of probabilistic or stochastic models of choice (Intriligator, 1973; Mattsson and Weibull, 2002; McFadden, 2005; Rieskamp, 2008; Blavatskyy and Pogrebna, 2010; Ryan, 2018). The degree of choice inconsistency may reflect individual differences in uncertainty about an adopted choice strategy, which in turn can determine the degree of emotional involvement when performing a financial choice task. In a typical risky decision, one option (the risky option) has multiple (for example, two) probabilistic outcomes (for example, 50-50%) while the other option (the safe option) provides an outcome with certainty. Various characteristics of choice options, such as the expected value (EV; i.e., the summed probability-weighted returns for each option) and inherent risk (i.e., the range of possible outcomes) affect the subjective value of options (Tobler et al., 2007). According to the traditional view, these features are represented and combined into options, which are then evaluated in a context-specific adaptive process (Louie et al., 2013, 2015).

Neuroscientific methods have provided insights into the brain processes and structures relevant for value-based decision making, and recent work supports the hypothesis that choice may be stochastic (Webb et al., 2019; Kurtz-David et al., 2019). In risk-averse participants, blood oxygen level dependent (BOLD) activity in anterior cingulate cortex (ACC) increases with risky choice probability (Christopoulos et al., 2009), whereas risk perception has been associated with activity in the insula (Bossaerts, 2010). In contrast, increased activity to safer options in the lateral prefrontal cortex (lPFC) is interpreted to function as a “safety” signal, playing an inhibitory role (Christopoulos et al., 2009). Moreover, the subjective value of risky options correlates with activity in the lPFC (Tobler et al., 2007, 2009; Holper et al., 2014). This link contributes causally to risk-dependent decisions (Knoch et al., 2006; Fecteau et al., 2007). Thus, differences in risk-related behavior appear to result from the interplay between several brain areas. However, relatively little is known about the temporal dynamics of these activations.

Decision making can be divided into multiple phases (Rangel et al., 2008): 1) a perception phase, where information is processed and integrated (e.g., identification and representation of stimulus features and available actions); 2) an option appraisal phase (e.g., assigning value to possible options); 3) a decision phase (option selection informed by valuation); 4) a response phase (motor action) and 5) and an outcome appraisal or feedback evaluation phase. Most event-related potentials (ERPs) studies in risky decision making have focused on evaluating outcomes (Polezzi et al., 2010; Schuermann et al., 2012; Zhang et al., 2014; Zheng et al., 2015; Xu et al., 2016; Fernandes et al., 2018) or anticipating outcomes (Zheng et al., 2015, 2020). Evidence from ERP studies suggests that assessment of expected value includes both early processes (100-200 ms after stimulus onset, more automatic and related to sensation) and late processes (450-650 ms after stimulus onset, more related to cognitive control). Both are believed to influence choice behavior (Bargh and Chartrand, 1999; Harris et al., 2013; Gu et al., 2018). If behavioral differences between participants result from the adoption of automatized, stereotypical heuristics, they may involve the use of highly trained mathematical calculations that offer quick access to solutions stored in long-term memory (see Logan’s instance theory; Logan, 1988). This rapid access to memory can be reflected in differences in neural activity that should be noticeable earlier in time than if they depended on subsequent controlled processes (in Logan’s terminology, algorithmic-based processing).

Several stimulus-locked ERP components have been identified during perception, evaluation and decision phases that follow the presentation of options and precede the participant’s response. These include the parietal N1 component (Eimer, 1998; Hillyard et al., 1998), also called N170 (Emmorey et al., 2017), N180 or N2, as an index of attention. The P200, with a latency of about 200 ms after stimulus onset, is supposed to reflect automatized stimulus evaluation (Luck and Hillyard, 1994; Testa et al., 2020), including rapid access to long-term memory representations, and therefore may be related to choice consistency across trials. ERP components that have been analyzed with regard to risk-taking include the medial frontal negativity (MFN, or frontocentral N2), a negativity peaking 250-300 ms after stimulus onset (Gehring and Willoughby, 2002) followed by the mid-latency component P3 (P300). The stimulus-related MFN may be part of a family of ERP components that includes the feedback-related negativity (FRN) and the error-related negativity (ERN) (Chandrakumar et al., 2018), which are elicited particularly in conditions in which stimulus or response conflict is present. These components are all thought to derive from the activation of the ACC, which may be associated with the perceived need to exert cognitive control in response to uncertain stimulus/contextual conditions (Botvinick et al., 2001). The P300 component is thought to reflect the end of the evaluation process (Coles et al., 1985), and its amplitude may reflect subjective uncertainty (Donchin and Coles, 1988). When evoked during outcome appraisal, the P300 has been associated with the motivational significance of the outcome (Nieuwenhuis et al., 2005), an association enhanced by risk-taking behavior (Polezzi et al., 2010). However, the P300 is sometimes difficult to distinguish from a late positive potential (LPP), which has been associated with the processing of emotional stimuli (see Hajcak et al., 2009) and with controlled processes involved in the modulation of subjective value (Harris et al., 2013).

The purpose of this study is to capture individual differences in risky decision making and to investigate the brain dynamics associated with such differences. We analyzed the stimulus-locked ERPs that follow the presentation of risky choice options and precede the participant’s responses. Specifically, we focused on the perception (early ERPs), evaluation (mid-latency ERPs) and decision (late ERPs) processes elicited by risky choice options. We expect the degree of choice consistency to be reflected in the ERP components associated with automatized stimulus processing, such as the anterior P200. We also anticipate that the MFN will play a role in the evaluation processes and that its amplitude will index the use of controlled processing and thus be associated with non-automated, less consistent and individual differences in behavior. We expect P300 to be associated with option selection, an association that could be modulated by risk aversion, as the motivational significance of a risky choice can vary between individuals with different attitudes towards risk. Finally, we expect LPP to be related to late modulation of value signals that should vary between individuals with different attitudes towards risk.

## 2 Methods

### 2.1 Participants

Healthy adults were recruited through advertisements posted on local community hubs. Prior to participation, they were informed about procedures and provided written informed consent for their participation in line with the Declaration of Helsinki (World Medical Association., 2001) and the protocol approved by the corresponding Swiss authority (Cantonal Ethic Research Committee of Geneva (CCER #2019-00095). All participants were screened in order to exclude any history of psychiatric, neurological or sleep disorders (Buysse et al., 1989; Spitzer et al., 2006; Kroenke et al., 2001). The final sample comprised N=24 participants (women =12, mean age = 38.6, SD = 10.5, range 27-61, 21 right-handed). All had normal or corrected-to-normal vision and all were naive to the decision-making task.

### 2.2 Risky decision task

We used a risky decision task similar to protocols previously described elsewhere (Tobler et al., 2007, 2009; Christopoulos et al., 2009; Holper et al., 2014, 2017). The task consisted of 20 blocks of 20 trials each. Participants received written task instructions explaining that on each trial they had to choose between two options, a safe (i.e., risk-free) option and a risky (i.e., variable) option. Each option comprised two outcomes. In the safe option the two outcomes were identical (the safe value SO). The risky option always consisted of two divergent outcomes, one higher and one lower than SO. For both options, participants were told that each outcome of the pair had a 50/50 chance of being drawn. The mean of the divergent outcomes for risky options defined the expected value (EV). The difference between the two outcomes defined the *risk level*. We used four risky options differentiated by two risk levels (*Risk_low_*, *Risk_high_*) and two EVs. For EV = 30 (i.e., EV_*small*_), the pair of outcomes for the risky option were 15 and 45 (*Risk_low_*) or 10 and 50 (*Risk_high_*). For EV = 60 (i.e., EV*_large_*), the outcomes were 40 and 80 (*Risk_low_*) or 30 and 90 (*Risk_high_*). Selecting the risky option resulted in a random draw (i.e., with the same probability 50%) of getting the low or high outcome, regardless of its amount.

Selecting the risk-free option the random draw always resulted in the SO. For each risky option, the displayed SO was randomly selected in a set of outcomes distributed within a range bounded by the two alternatives of the risky option according to a pseudo-normal distribution, centered on the average certainty equivalents, identified in previous research (Holper et al., 2017). Hence, four series of *n* = 100 discrete outcomes of SO following pseudo-normal distributions were generated by rounding the floating-point values of the corresponding normal distribution 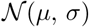, with mean *μ* and standard deviation *σ*. We set 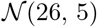 for EV_*small*_ and *Risk_low_*, 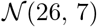 for EV_*small*_ and *Risk_high_*, 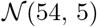 for EV_*large*_ and *Risk_low_*, and 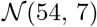 for EV_*large*_ and *Risk_high_*.

Participants were advised that at the end of the experiment one trial out of all 400 trials would be randomly selected and that they would receive 10% of the outcome determined according to the probability of the option the participant chose in the selected trial.

### 2.3 Behavioral procedure

Participants sat in a comfortable chair and received written task instructions. The experiment was run with the lights off. The background of the computer screen was dark gray. Figure 1 illustrates the schematic layout of a trial. Each trial began with participants pressing the space bar. Next, they were asked to fixate a white cross in the center of a computer screen, placed 65 cm away, for a random interval lasting 1000, 1200, 1400, 1600, 1800, or 2000 ms. Options were presented to the left and right of the fixation cross with a lateral visual angle of ~5.3° and vertical visual angle of ~3.7°. The sides at which the safe and risky options were presented, as well as the vertical placement of the higher outcome of the risky option (i.e., top or bottom) were randomly selected. To minimize learning, no feedback was provided regarding the chosen option. However, the fixation cross turned purple when a choice was made. The inter-trial interval (i.e., 500, 700, 900, 1100, 1300 or 1500 ms) was randomly selected to avoid cross- and within-trial contamination of EEG signals.

**Figure 1:**
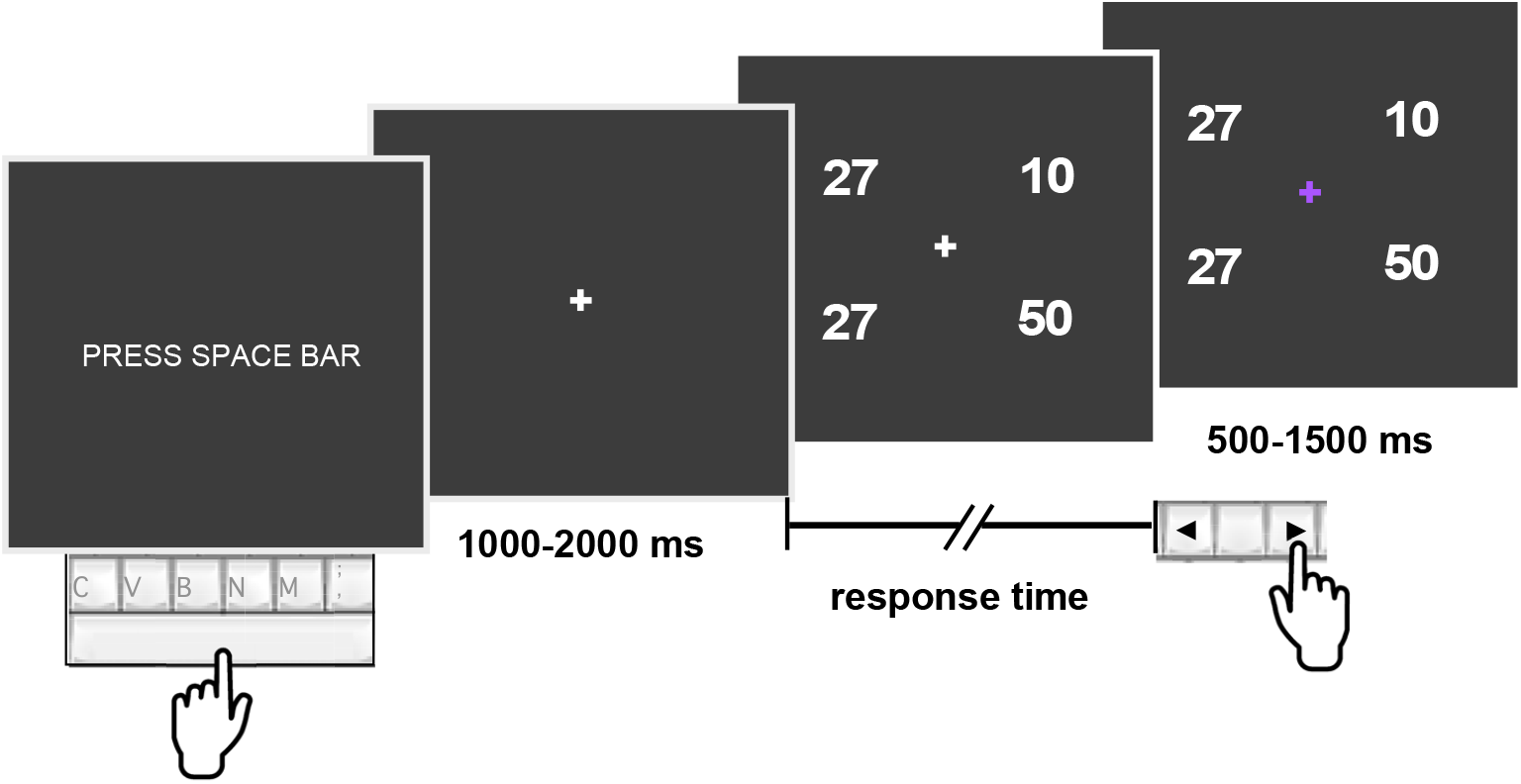
Schematic representation of a trial. Each trial started when the participant pressed the spacebar. A fixation cross was presented for 1-2 s, followed by the presentation of two choice options. Participants conveyed their choice by pressing the left or right arrow key on the computer keyboard with the index finger of their right hand. In this example, the risk-free (safe) option is presented on the left side, which means that choosing the left arrow key always results in 27. The risky option (in this example on the right side) always consisted of two divergent alternatives, one higher and one lower than the safe value. In this case, by choosing the right arrow key 10 or 50 are randomly drawn with equal probability, irrespective of their position (top or bottom) on the screen. The fixation cross turned purple after the choice was made and no feedback was provided on the outcome of the random draw.

### 2.4 Behavioral measurements

A preprocessing step consisted in the analysis of response times (RTs) for each participant and each condition. To avoid the effect of exceedingly long RTs on the analysis of behavioral performance, we excluded the trials with RT beyond the 98th percentile of the corresponding condition. Following previous research (Christopoulos et al., 2009), we estimated the attitude towards risk from the choices made in the task. Specifically, for each option we calculated the certainty equivalent (CE), which is the theoretical amount at which the participant is equally likely to choose the safe or the risky option (Luce et al., 1999). For each trial, we set a binary variable *safe choice* to 0 if the participant rejected the safe option (i.e., selected the risky option) and to 1 when the safe option was selected. For each trial, the proportion of *safe choices* is plotted as a function of the corresponding safe option SO (see Figure 2). For each of the two risk levels the data was fitted with the following pseudo sigmoid function (Strasburger, 2001):

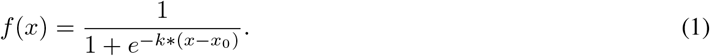

where *x* takes only values within the range of risky option outcomes, *x*_0_ is the minimum outcome and *k* is the slope parameter of the sigmoid function (Figure 2). For each condition, the analysis of the S-curves determined the cEs, corresponding to the points on the x-axis for which *f*(*x*) = 0.5 (green arrow in Figure 2). The nonlinear least-squares estimates of these parameters were obtained using the R package `nls2` (Grothendieck, 2013).

**Figure 2:**
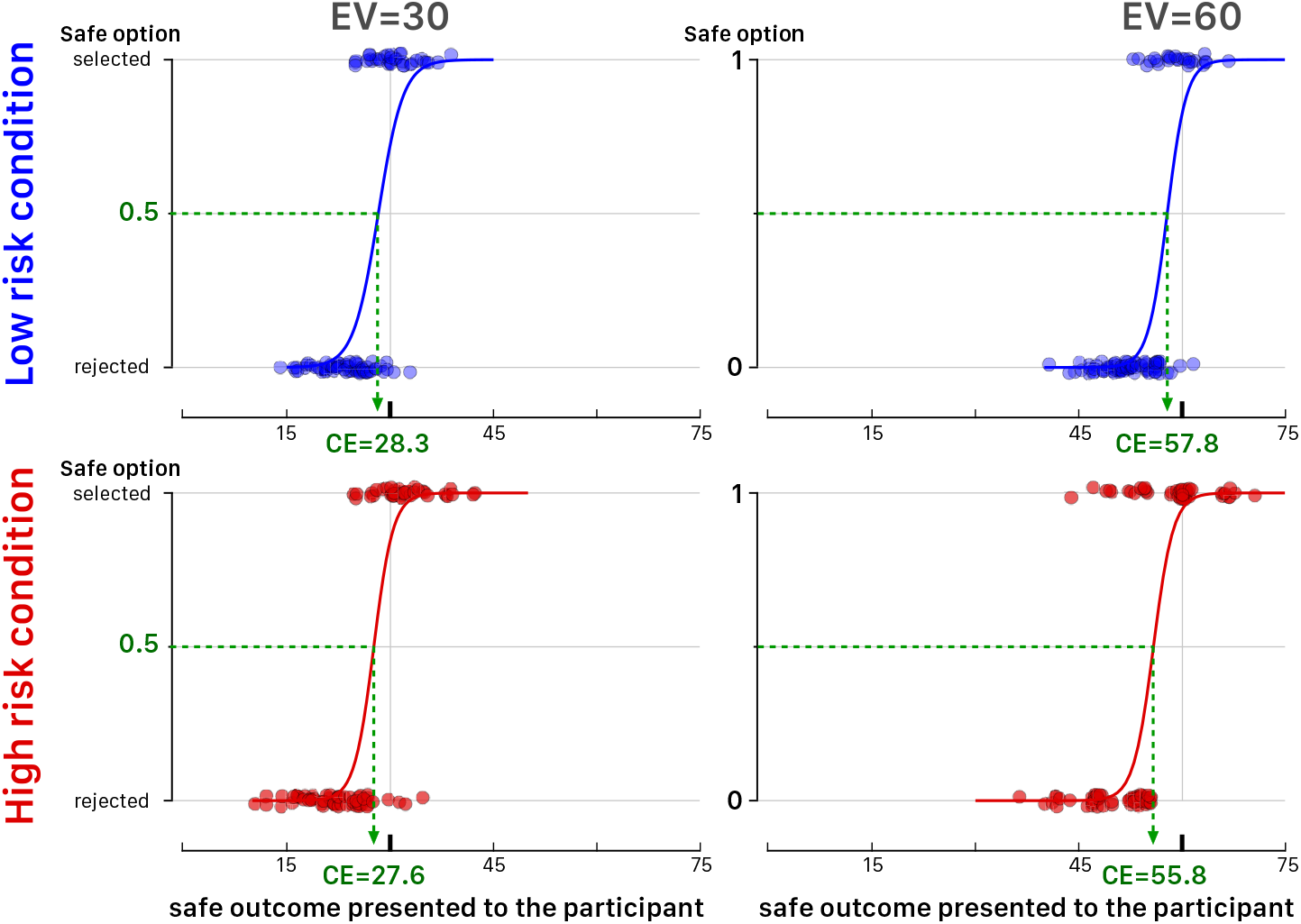
Example behavior for one participant (2Ba7) during the four conditions of the task. On each trial the participant could select the safe option (binary value 1 on the y-axis) or reject it (binary value 0 on the y-axis), corresponding to selection of the risky option. Each trial is represented by one dot. The top and bottom panels illustrate choices in the low (blue dots) and high (red dots) risk conditions, respectively. The left and right panels illustrate choices for the small (EV=30) and large (EV=60) expected values, respectively. The x-axis represents the safe outcome presented at each trial. The sigmoid curves allow determining the risk attitude of the participant as a function of the safe outcome and estimate the *certainty equivalent* (CE).

Risk attitude was defined by the difference between the CEs of the high- and the low-risk conditions, keeping EV constant (Christopoulos et al., 2009). Thus, larger positive (or negative) differences correspond to increasing risk aversion (or risk proneness). For the two levels of EV (i.e., EV=30 and EV=60) and for each participant P, we computed the corresponding *risk aversion* parameters (i.e., *RA*_small(P)_ and *RA*_large(P)_, respectively). Both values were standardized to EV=30 to be averaged and obtain *RA*_mean(P)_. The product *RA*_(P)_ = *RA*_small(P)_ × *RA*_large(P)_ and the value *RA*_mean(P)_ are good descriptors of individual risk aversion and for categorization of participants into groups. A participant P was categorized in the group: *(i) Risk-seeking*, if both values *RA*_small(P)_ and *RA*_large(P)_ were less than −1.0. or if *RA*_(P)_ < −1.0; *(ii) Risk-neutral*, if −1.0 < *RA*_(P)_ < 1.0; *(iii) Risk-averse*, if *RA*_(P)_ > 1.0. The cutoff values −1.0 and 1.0 were based on a preliminary analysis of the distribution of *RA*_(P)_ considering a Gaussian mixture model with three groups of participants (i.e., *Risk-seeking, Risk-neutral, Risk-averse*).

It appears reasonable to assume that risk attitudes should be similar across the conditions with small and large expected values. By extension, any participant not included in the previous groups and showing a large incongruity in risk attitude between EV=30 and EV=60 (e.g., showing risk aversion at one value and strong risk-seeking attitude at the other value) is categorized as *Risk-incongruent*. We set such risk-incongruity with the criterion | *RA*_large(P)_ / *RA*_small(P)_ | > 4.0 or | *RA*_large(P)_/*RA*_small(P)_ | < 0.25.

At the end of the session, for each participant P and for each condition j, we also computed a measure of within-participant consistency: *consistency_j_* = (*n*_1*j*_ + *n*_0*j*_)/*N*_tot*j*_, where *n*_1*j*_ is the count of any trial *i* during condition *j* when the participant selected the risk-free option given SO_*ij*_ > CE_*j*_, *n*_0*j*_ is the count of any trial *i* during condition *j* when the participant selected the risky option given SO_*ij*_ < CE_*j*_, and *N*_tot*j*_ is the overall number of trials in condition *j*. A consistency equal to 1 corresponds to a S-curve modified into a Heaviside step function. We used the average consistency across the four experimental conditions *c*_mean(P)_ to characterize each participant P.

### 2.5 Electrophysiological recordings

EEG signals were recorded continuously at a 1024 Hz sampling frequency (24 bit resolution) with an ActiveTwo MARK II Biosemi EEG System (BioSemi B.V., Amsterdam, The Netherlands) using NeuroSpec Quick Caps with 64 scalp Ag/AgCl active electrodes (extended 10/20 layout). Two pairs of bipolar electrodes were used to record eye movements. The EOG electrodes were placed at the outer canthi of both eyes and above and below the left eye, to record horizontal and vertical eye movements, respectively. EEG and EOG were recorded with a band-pass filter of 0.05-200 Hz. Impedance was kept below 5 KΩ. Scalp electrodes were referenced to the left mastoid during recording and re-referenced off-line to the average of the left and right mastoid.

### 2.6 EEG data analysis

EEG data were preprocessed and analyzed with the ERPLAB toolbox (Lopez-Calderon and Luck, 2014) and with dedicated MATLAB scripts (MATLAB, The MathWorks, Inc). EEG data were segmented into epochs of 800 ms (baseline = 160 ms) based on markers identifying the start of the stimulus, i.e., the presentation of the options. Trials with RT<640 ms, i.e., shorter than the ERP response window, were excluded to minimize contamination by response-induced activity. Trials with RT>6000 ms and trials with RT beyond the 99 percentile were also excluded from further analysis (mean rejected trials = 0.052, SD = 0.047). Ocular artifacts (blinks and saccades) were removed from the signals with the eye movement correction procedure (EMCP) (Gratton et al., 1983). In addition, epochs were visually inspected for contamination by muscular and recording artifacts and excluded from further analyses if such artifacts were detected. Epochs passing all the exclusion criteria (mean = 82%) were low-pass filtered at 40 Hz. Average ERP waveforms were derived for each participant, electrode and condition.

As mentioned in the introduction, here we focused on the perception and valuation phases of the decision process. We defined 3 sets of frontocentral electrodes (medial: Fz, F1, F2, FCz, FC1, FC2; left: F3, F5, F7, FC3, FC5, FT7; right: F4, F6, F8, FC4, FC6, FT8) and 3 sets of centro-parietal electrodes (medial: Pz, P1, P2, CPz, CP1, CP2; left: P3, P5, P7, CP3, CP5, TP7; right: P4, P6, P8, CP4, CP6, TP8). Poor ERP signals from electrodes included in the sets were identified using the average standardized measurement error (SME) (Luck et al., 2021). Electrodes with an average SME > 3 microvolts (*μ*V) were excluded.

## 3 Results

### 3.1 Behavior

#### 3.1.1 Effect of Risk Level

Each of the 24 participants performed 400 trials for a total of 9600 trials. Out of these, 37 trials with exceedingly long RTs were eliminated during preprocessing, leaving 9563 trials for the behavioral analysis. RTs of all participants were grouped together and we analyzed the median RTs as a function of the safe outcome for all the outcomes with more than 5 valid trials (Figure 3A). In all four conditions, RTs followed inverted-U shaped curves, suggesting that decision making took more time when the safe values were close to the behavioral indifference points (certainty equivalents). We used Aikake Information Criterion (AIC) model selection to distinguish polynomial models of 2nd and 3rd order describing the relationship between median RTs and safe outcomes at all conditions. Third order models showed higher fits than 2nd order models, as well as larger adjusted *R*^2^ and lesser Leave-one-out cross-validation (LOOCV) prediction error (*F*(3, 15) = 9.109, *p* < 0.01, *R*^2^ = 0.5747, LOOCV= 78482 for [EV=30, *Risk_low_*]; *F*(3, 12) = 5.103, *p* < 0.05, *R*^2^ = 0.4507, LOOCV= 96574 for [EV=30, *Risk_high_*]; *F*(3, 11) = 3.937, *p* < 0.05, *R*^2^ = 0.3862, LOOCV= 96385 for [EV=60, *Risk_low_*]; and *F*(3, 12) = 18.31, *p* < 0.001, *R*^2^ = 0.7759, LOOCV= 20707 for [EV=60, *Risk_high_*]). Thus, as would be expected, more difficult choices took longer, indicating that participants did not respond randomly in our task.

**Figure 3:**
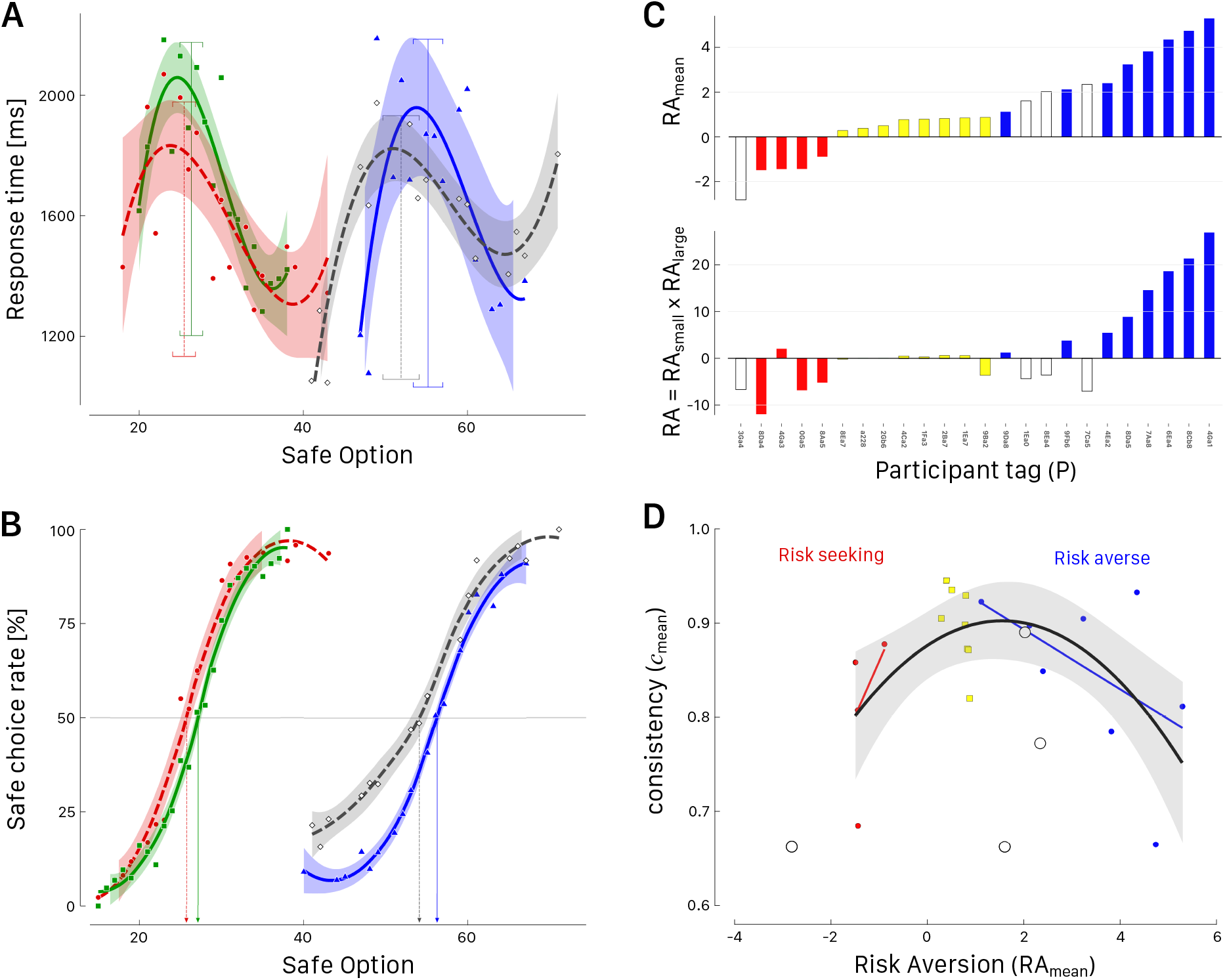
Behavioral results. **A.** Median response times as a function of the safe outcomes across the four conditions (green squares: EV*_small_* and *Risk_low_*; red dots: EV*_small_* and *Risk_high_*; blue triangles: EV*_large_* and *Risk_low_*; empty grey diamonds: EV*_large_* and *Risk_high_*). All curves correspond to 3rd order polynomial fit and shaded areas to 95% confidence intervals. Vertical lines and 95% error bars indicate the values of the CEs for each condition. **B.** Safe choice rate as a function of proposed safe outcome in the four conditions of the risky decision task. Dashed curves correspond to high risk conditions. All curves are smoothed by locally weighted quadratic regression (loess) with closest 4/5 of the total data points and tricube weighting. Shaded areas correspond to 95% confidence intervals. **C.** Risk aversion parameters (*RA*_mean_ and *RA*) for each participant identified by a unique tag P. Participants are ordered by *RA*_mean_ and categorized as *Risk-averse* (blue bars), *Risk-neutral* (yellow bars), *Risk-seeking* (red bars), and *Risk-incongruent* (empty bars). **D.** Consistency (*c*_mean_) as a function of risk aversion (*RA*_mean_). Blue dots correspond to *Risk-averse*, yellow squares to *Risk-neutral*, red dots to *Risk-seeking*, and empty dots to *Risk-incongruent* participants. A black curve shows the quadratic fit, for all values except the *Risk-incongruent* participants, and its corresponding 95% confidence limits (shaded area). Linear fits for *Risk-averse* (blue line) and *Risk-seeking* participants (red line) are also indicated.

To determine individual risk attitudes, we estimated CEs for all participants in the four conditions, as illustrated for one participant in Figure 2. In line with overall risk aversion, the average CEs were smaller than the corresponding EVs and within given EVs, high-risk conditions resulted in lower CE than low-risk conditions. For instance, we observed CE_1_ = 26.35±0.71 for [EV=30, *Risk_low_*], CE_2_ = 25.47±0.71 for [EV=30, *Risk_high_*], CE_3_ = 55.24±0.92 for [EV=60, *Risk_low_*] and CE_4_ = 51.95±1.13 for [EV=60, *Risk_high_*]. These values were close to the certainty equivalents CE estimated from the curves of the safe choice rate as a function of the safe outcomes (Figure 3B) when all participants were grouped together, namely 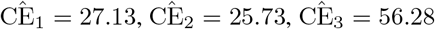 and 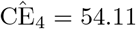. Thus, irrespective of the exact method used, participants on average were risk averse in our task.

#### 3.1.2 Categorized groups of participants

The participants were categorized as *Risk-averse* (RA), *Risk-neutral* (RN), *Risk-seeking* (RS) or *Risk-incongruent* following the assessment of their risk attitude with the criteria described in the Methods section. The groups (Table 1, Figure 3C) were well characterized by *RA*_mean_ (one-way ANOVA, *F*(3, 20) = 13.581, *p* < .001, and a large effect size 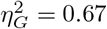). At face value (Figure 3C), it could be argued that one *Risk-incongruent* participant should be included in the *Risk-seeking* group and a couple of *Risk-incongruent* participants might be included in the *Risk-averse* group. However, we kept the initial criteria for categorization with the aim to clearly separate participants with congruent risk attitude from participants with incongruent risk attitude and thereby achieve as much group homogeneity as possible for the analysis of ERPs.

**Table 1:** Descriptive statistics of behavioral analysis for the three groups of participants. Median (mean ± SEM).

| Behavioral group | Risk Aversion |  |  |  | Consistency |
| --- | --- | --- | --- | --- | --- |
| | $RA_{\text{small}}$ | $RA_{\text{large}}$ | $RA_{\text{mean}}$ | $RA$ | $c_{\text{mean}}$ |
| **Risk-averse** ($N = 8$) | 3.38<br>3.75±0.63 | 3.75<br>3.02±0.50 | 3.53<br>3.38±0.51 | 11.72<br>12.60±3.25 | 0.87<br>0.85±0.03 |
| **Risk-neutral** ($N = 8$) | 0.74<br>0.94±0.34 | 0.66<br>0.40±0.28 | 0.79<br>0.67±0.08 | 0.17<br>-0.21±0.51 | 0.90<br>0.90±0.01 |
| **Risk-seeking** ($N = 4$) | -2.29<br>-1.68±1.47 | -0.04<br>-0.94±1.62 | -1.44<br>-1.31±0.14 | -6.02<br>-5.49±2.89 | 0.83<br>0.81±0.04 |
| **Risk-incongruent** ($N = 4$) | -1.11<br>-2.41±1.41 | 4.52<br>3.98±1.05 | 1.82<br>0.79±1.21 | -5.52<br>-5.41±0.86 | 0.72<br>0.75±0.05 |

For *RA* only, the Levene’s Test for homogeneity of variance was significant (*F*(3, 20) = 8.7668, *p* < .001). Therefore, we compared the groups with robust statistics (R package WRS2; Mair and Wilcox, 2020) and found that *Risk-averse* vs. *Risk-neutral* and *Risk-neutral* vs. *Risk-seeking* differences were not significant (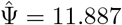, *p* = .071 and 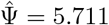, *p* = .285, respectively). It is interesting to note that the categorization based on risk attitude (Table 1) showed a significant main effect of *Behavioral group* for consistency (*c*_mean_, *F*(3, 20) = 3.427, *p* = .037, 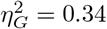). This result suggests a possible relationship between consistency and (categories of) risk aversion and we focus on this relationship next.

#### 3.1.3 Consistency and risk aversion

The relationship between consistency and risk aversion, excluding the *Risk-incongruent* participants could be fitted by a quadratic curve (black curve in Figure 3D) with statistic *F*(2, 17) = 4.783, *p* < 0.05. Overall, these results (Figure 3D) suggest that in our task, the group of participants categorized as *Risk-neutral* was characterized by the highest ratio of consistency (see also Table 1). In other words, the greater the tendency of participants to seek or avoid risk, the less consistently they behaved in our task.

### 3.2 Grand average ERPs

The main focus of this paper is the investigation of the relationship between individual differences in risky decision making and the brain dynamics occurring while making these choices, as identified using ERPs. Because of its small size, we did not analyze the averaged ERPs of the RS group. Grand average ERP waveforms from the six electrode sets for RN and RA groups are illustrated in Figure 4. The frontal P200 (200-300 ms) and the MFN (300-400 ms) were analyzed at frontocentral sites, while the parietal N100 (N1; 150-250 ms), P300 (275-425 ms) and LPP (450-600 ms) were analyzed at centro-parietal sites.

**Figure 4:**
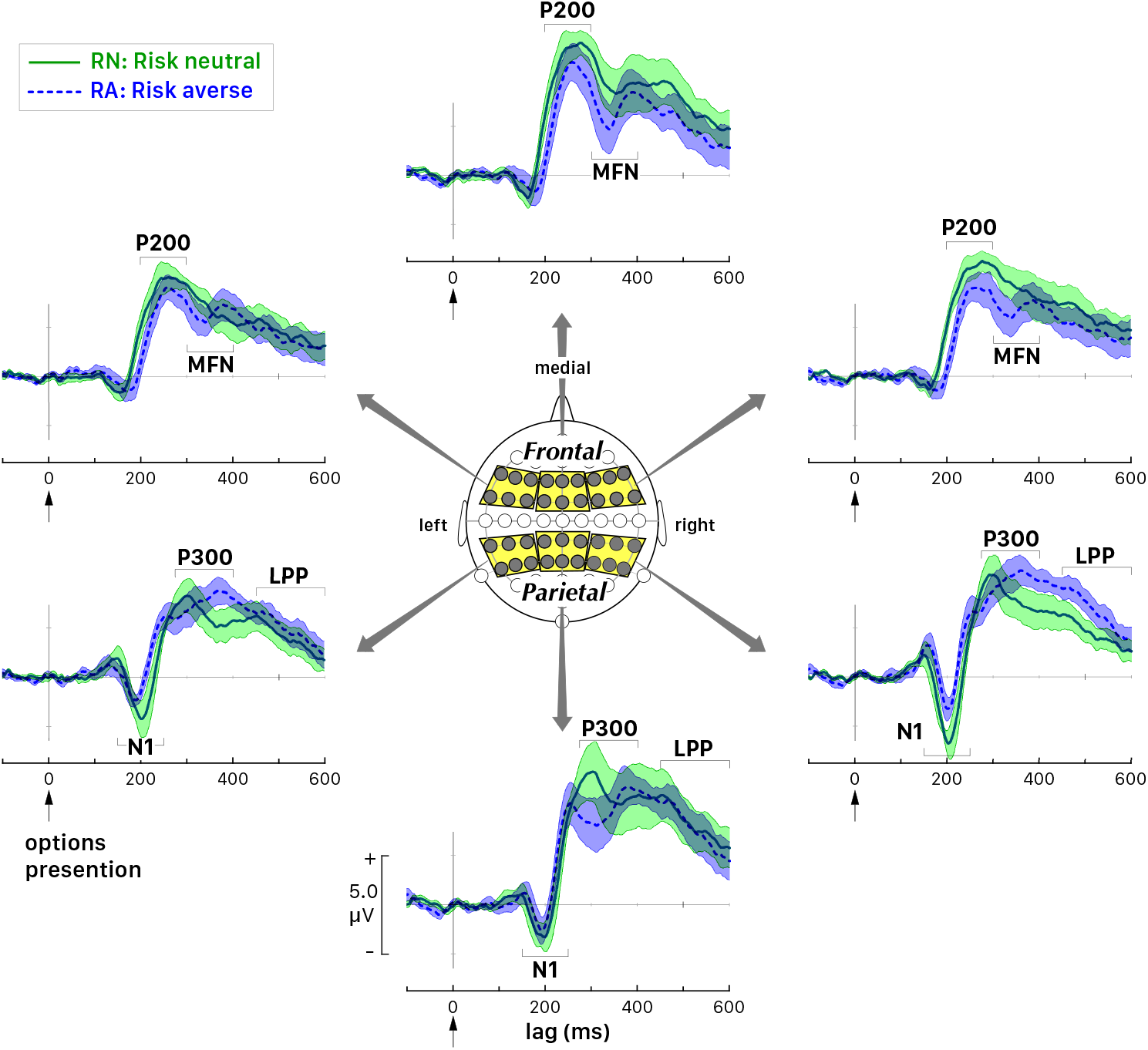
Distribution of cortical activity related to option presentation for participants classified as *Risk-neutral* (RN, green solid lines) and *Risk-averse* (RA, blue dashed lines). Mean activity (grand average ERPs) and confidence intervals (± SEM, shaded areas) for three sets of frontocentral and three sets of centro-parietal electrodes. MFN: medial frontal negativity. LPP: late positive potential.

The brain processes associated with the actual choice of the uncertain (risky) and certain (safe) option on each trial were investigated by sorting trials according to the decision made by the participants. For this analysis, we pooled the four conditions (EV=30/*Risk_low_*, EV=30/*Risk_high_*, EV=60/*Risk_low_*, EV=60/*Risk_high_*) for the corresponding grand average waveforms of all participants (N=20, excluding the four participants classified as *Risk-incongruent*) at the central and lateral electrode sets (Figure 5A).

**Figure 5:**
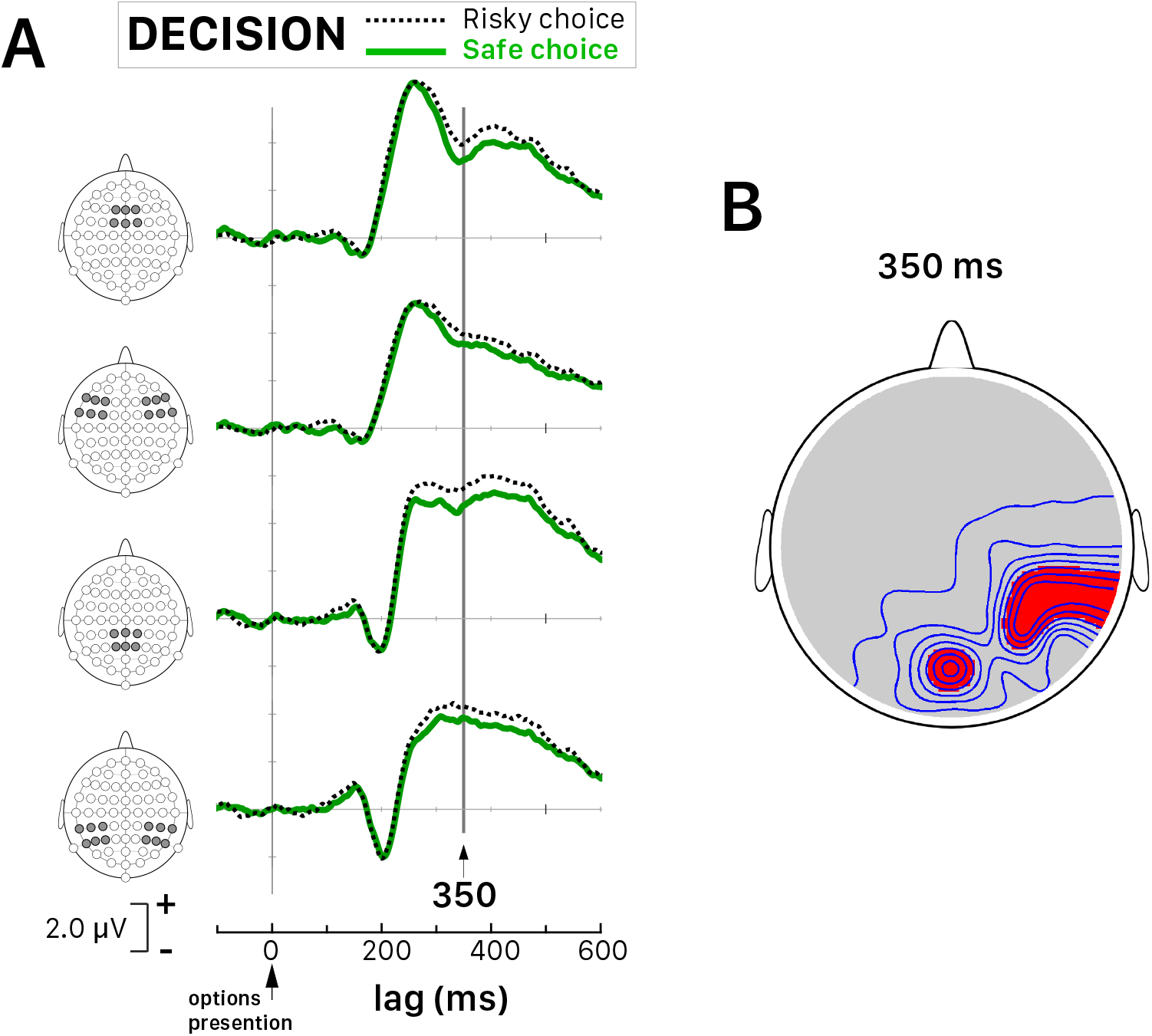
Distribution of cortical activity related to decision. **A.** Grand average ERP waveforms (20 participants), sorted by subsequent selection of risky or safe choice, at medial and lateral electrode sets (merging left and right sets). The latency of 350 ms (black vertical line) corresponds to the lag after stimulus onset with the statistically largest difference between selecting risky versus safe options. **B.** Topographic map of the significant difference between risky and safe choices at 350 ms (*red* = *p* < .05; *grey* = *p* > .05).

Paired *t*-tests on the difference between risky and safe choices were computed for individual average waveforms at each of the 64 electrode sites to determine at which latency these waveforms differed, if any. Bonferroni-Holm correction was performed on the 64 electrode sites in order to control for multiple comparisons (Holm, 1979). We found a greater positivity at 350 ms after stimulus onset at several right and mid-parietal electrode sites when participants chose the risky option. Mapping the significant differences between risky and safe choices at this latency (Figure 5B) showed that brain activity at right parietal electrode sites differed depending on the decision, in line with the hypothesis that mid-latency evoked activity (P300) is associated with behaviorally relevant information and option selection. Moreover, these neural findings reinforce the conclusion based on the behavioral data that participants were not choosing randomly.

To investigate the effects of risk attitude, we compared RA and RN groups, first by examining the components’ mean amplitude. A two-way (Group × Electrode-set-position) ANOVA was performed for each ERP component (Table 2). We observed a significant main effect of Group for the parietal N1 and the frontal P200 (N1: *F*_(1, 42)_ = 7.39, *p* < .01, *η*^2^ =0.15; P200: *F*_(1, 42)_ = 6.12, *p* < .05, *η*^2^ = 0.13). No other significant main effect was found, although there was a trend for a main effect of Electrode-set-position for LPP (*F*_(1, 42)_ = 3.14, *p* = .05, η^2^ = 0.13). No significant interaction effect was found. Thus, risk averse participants processed the selection of risky choice options differently than risk neutral participants.

**Table 2:**
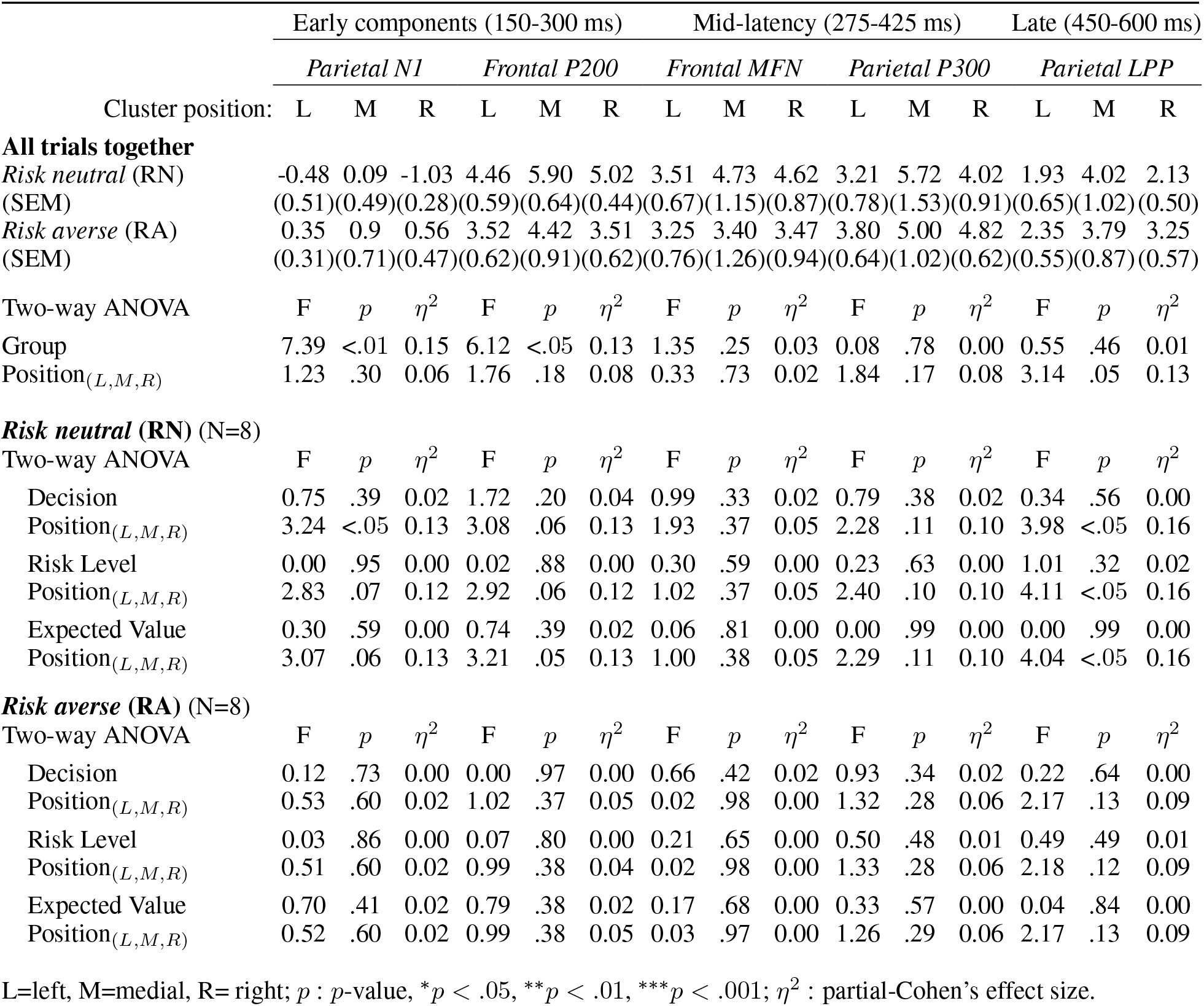
Descriptive statistics of ERP components’ mean amplitude (*μ*V) for each group at all positions, regardless of the risk level and expected value and ANOVA results for each factor separately.

|  | Early components (150-300 ms) |  |  |  |  |  | Mid-latency (275-425 ms) |  |  |  |  |  | Late (450-600 ms) |  |  |  |
| --- | --- | --- | --- | --- | --- | --- | --- | --- | --- | --- | --- | --- | --- | --- | --- | --- |
|  | *Parietal N1* |  |  | *Frontal P200* |  |  | *Frontal MFN* |  |  | *Parietal P300* |  |  | *Parietal LPP* |  |  |  |
| Cluster position: | L | M | R | L | M | R | L | M | R | L | M | R | L | M | R |  |
| **All trials together** |  |  |  |  |  |  |  |  |  |  |  |  |  |  |  |  |
| *Risk neutral* (RN) | -0.48 | 0.09 | -1.03 | 4.46 | 5.90 | 5.02 | 3.51 | 4.73 | 4.62 | 3.21 | 5.72 | 4.02 | 1.93 | 4.02 | 2.13 |  |
| (SEM) | (0.51) | (0.49) | (0.28) | (0.59) | (0.64) | (0.44) | (0.67) | (1.15) | (0.87) | (0.78) | (1.53) | (0.91) | (0.65) | (1.02) | (0.50) |  |
| *Risk averse* (RA) | 0.35 | 0.9 | 0.56 | 3.52 | 4.42 | 3.51 | 3.25 | 3.40 | 3.47 | 3.80 | 5.00 | 4.82 | 2.35 | 3.79 | 3.25 |  |
| (SEM) | (0.31) | (0.71) | (0.47) | (0.62) | (0.91) | (0.62) | (0.76) | (1.26) | (0.94) | (0.64) | (1.02) | (0.62) | (0.55) | (0.87) | (0.57) |  |
| Two-way ANOVA | F | p | $\eta^2$ | F | p | $\eta^2$ | F | p | $\eta^2$ | F | p | $\eta^2$ | F | p | $\eta^2$ | |
| Group | 7.39 | <.01 | 0.15 | 6.12 | <.05 | 0.13 | 1.35 | .25 | 0.03 | 0.08 | .78 | 0.00 | 0.55 | .46 | 0.01 |  |
| Position<sub>(L,M,R)</sub> | 1.23 | .30 | 0.06 | 1.76 | .18 | 0.08 | 0.33 | .73 | 0.02 | 1.84 | .17 | 0.08 | 3.14 | .05 | 0.13 |  |
| ***Risk neutral* (RN) (N=8)** |  |  |  |  |  |  |  |  |  |  |  |  |  |  |  |  |
| Two-way ANOVA | F | p | $\eta^2$ | F | p | $\eta^2$ | F | p | $\eta^2$ | F | p | $\eta^2$ | F | p | $\eta^2$ | |
| Decision | 0.75 | .39 | 0.02 | 1.72 | .20 | 0.04 | 0.99 | .33 | 0.02 | 0.79 | .38 | 0.02 | 0.34 | .56 | 0.00 |  |
| Position<sub>(L,M,R)</sub> | 3.24 | <.05 | 0.13 | 3.08 | .06 | 0.13 | 1.93 | .37 | 0.05 | 2.28 | .11 | 0.10 | 3.98 | <.05 | 0.16 |  |
| Risk Level | 0.00 | .95 | 0.00 | 0.02 | .88 | 0.00 | 0.30 | .59 | 0.00 | 0.23 | .63 | 0.00 | 1.01 | .32 | 0.02 |  |
| Position<sub>(L,M,R)</sub> | 2.83 | .07 | 0.12 | 2.92 | .06 | 0.12 | 1.02 | .37 | 0.05 | 2.40 | .10 | 0.10 | 4.11 | <.05 | 0.16 |  |
| Expected Value | 0.30 | .59 | 0.00 | 0.74 | .39 | 0.02 | 0.06 | .81 | 0.00 | 0.00 | .99 | 0.00 | 0.00 | .99 | 0.00 |  |
| Position<sub>(L,M,R)</sub> | 3.07 | .06 | 0.13 | 3.21 | .05 | 0.13 | 1.00 | .38 | 0.05 | 2.29 | .11 | 0.10 | 4.04 | <.05 | 0.16 |  |
| ***Risk averse* (RA) (N=8)** |  |  |  |  |  |  |  |  |  |  |  |  |  |  |  |  |
| Two-way ANOVA | F | p | $\eta^2$ | F | p | $\eta^2$ | F | p | $\eta^2$ | F | p | $\eta^2$ | F | p | $\eta^2$ | |
| Decision | 0.12 | .73 | 0.00 | 0.00 | .97 | 0.00 | 0.66 | .42 | 0.02 | 0.93 | .34 | 0.02 | 0.22 | .64 | 0.00 |  |
| Position<sub>(L,M,R)</sub> | 0.53 | .60 | 0.02 | 1.02 | .37 | 0.05 | 0.02 | .98 | 0.00 | 1.32 | .28 | 0.06 | 2.17 | .13 | 0.09 |  |
| Risk Level | 0.03 | .86 | 0.00 | 0.07 | .80 | 0.00 | 0.21 | .65 | 0.00 | 0.50 | .48 | 0.01 | 0.49 | .49 | 0.01 |  |
| Position<sub>(L,M,R)</sub> | 0.51 | .60 | 0.02 | 0.99 | .38 | 0.04 | 0.02 | .98 | 0.00 | 1.33 | .28 | 0.06 | 2.18 | .12 | 0.09 |  |
| Expected Value | 0.70 | .41 | 0.02 | 0.79 | .38 | 0.02 | 0.17 | .68 | 0.00 | 0.33 | .57 | 0.00 | 0.04 | .84 | 0.00 |  |
| Position<sub>(L,M,R)</sub> | 0.52 | .60 | 0.02 | 0.99 | .38 | 0.05 | 0.03 | .97 | 0.00 | 1.26 | .29 | 0.06 | 2.17 | .13 | 0.09 |  |

L=left, M=medial, R= right; $p$ : $p$ -value, \* $p$ < .05, \*\* $p$ < .01, \*\*\* $p$ < .001; $\eta^2$ : partial-Cohen's effect size.

Secondly, within groups RA and RN, we aimed at identifying significant interactions between components and individual differences in behavior. For each participant, the mean amplitude of each ERP component was computed for each experimental condition at each of the three electrode set positions (left, medial and right; Table 2). We performed two-way (“Factor” × Electrode-set-position) ANOVAs for each ERP component, with 3 levels of Electrode-set-position (L, M, R) and “Factor” being either Decision (2 levels: *safe* or *risky* choice), Risk Level (2 levels: *Risk_high_*, *Risk_low_*), or Expected Value (2 levels: small EV=30, large EV=60). In risk neutral participants, we observed a significant main effect of Electrode-set-position with parietal N1 for the Decision factor (*F*_(2,42)_ = 3.24, *p* < .05, *η*^2^ = 0.13). Moreover, a significant main effect of Electrode-set-position was found for LPP with all factors (Decision: *F*_(2,42)_ = 3.98, *p* < .05, η^2^ = 0.16; Risk level: *F*_(2,42)_ = 4.11, *p* < .05, η^2^ = 0.16; Expected Value: *F*_(2,42)_ = 4.04, *p* < .05, η^2^ = 0.16). No other significant main effect and no significant interaction effect was found. These findings suggest that the processing of Decision, Risk Level and Expected Value was distributed across electrode positions.

### 3.3 Effect of stimulus condition in behavioral groups

Behavioral differences between the RN and RA groups suggest that they may be related to different topographic distributions and time courses of cortical activity for the factors Decision, Risk Level and Expected Value. As for the analyses above, we investigated the dynamics and scalp distribution of the cortical activity across the six electrode sets. For illustration purposes we show the analysis for factor Risk at the right parietal electrode set (Figure 6). These analyses were performed regardless of the decision taken (safe or risky option) and regardless of the expected value (EV=30 or EV=60). Within each group, we computed the differential curve between the ERP recorded during *Risk_high_* and *Risk_low_* and assessed its significance by paired *t*-test with *p* < .01 at each time bin (Figure 6B). We considered only intervals of consecutive significant bins lasting at least 5 ms. In the RN group, we observed that at the right parietal set *Risk_high_* evoked an activity which tended always to be larger than that evoked by *Risk_low_*, in particular in the interval 472-480 ms after stimulus onset. Conversely, in the RA group, the activity evoked by *Risk_high_* tended to be smaller than the activity evoked by *Risk_low_*, in particular in the interval 576-586 ms after stimulus onset (Figure 6C, D). Thus, a high level of risk evoked significantly less activity in the right parietal area of participants belonging to the RA group, compared to RN participants, supporting the hypothesis that risk affects late activity in a risk attitude-dependent fashion. In the remainder of this section, we describe these relationships for specific ERP components across all electrode sets, ordered by factor.

**Figure 6:**
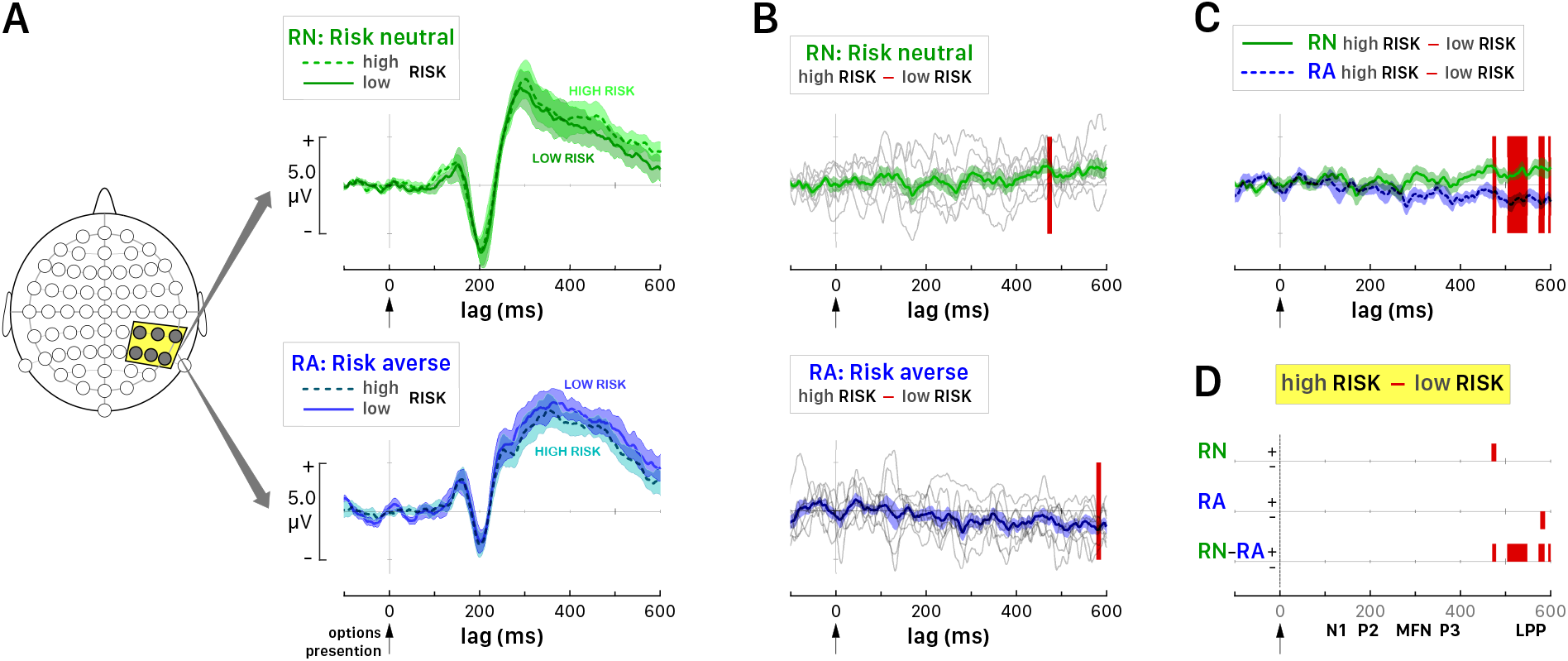
Activity evoked by different levels of risk at the right parietal set of six electrodes in *Risk neutral* (RN) and *Risk averse* (RA) participants. **A.** Grand average ERPs following risk level in RN (upper panel, green curves) and RA (lower panel, blue curves) groups. Solid and dark-colored lines show the averaged ERP for *Risk_low_* condition; dashed and light-colored lines correspond to *Risk_high_* condition. Shaded areas are delimited by ±SEM. **B.** Differential curves of averaged ERP evoked by *Risk_high_* minus averaged ERP evoked by *Risk_low_* for RN (green curve, upper panel) and RA (blue curve, lower panel). Shaded areas are delimited by ±SEM. The grey curves correspond to individual participants (N=8) in each group. The red labels correspond to the intervals with a level of significance *p* < .01 for comparison against amplitude zero. **C.** The differential curves of RN and RA groups with the respective confidence intervals as in panel B. The red labels indicate intervals where the differential curve in RN differed significantly (p < .01) from the differential curve in RA. **D.** Schematic plot showing the significant intervals of the differential curves corresponding to *Risk_high_* minus *Risk_low_* conditions for groups RN, RA and the difference RN minus RA. The red labels at the significant intervals point upward for positive and downward for negative values of the amplitude of the differential curve.

#### 3.3.1 Effect of Risk Level

Following the same procedure described above (Figure 6), we observed no significant effect of risk level over the left hemisphere (Figure 7). In the right hemisphere, we observed that in addition to the parietal electrode set (Figure 7) the effect of risk was also significant in the frontal set during LPP with the same dynamics observed at the parietal area. Compared to right parietal regions, right frontal regions tended to show a stronger negative effect on the amplitude of the differential curve for RA and a weaker positive effect for RN. More importantly, in both regions, relatively late risk processing depended on risk attitude.

**Figure 7:**
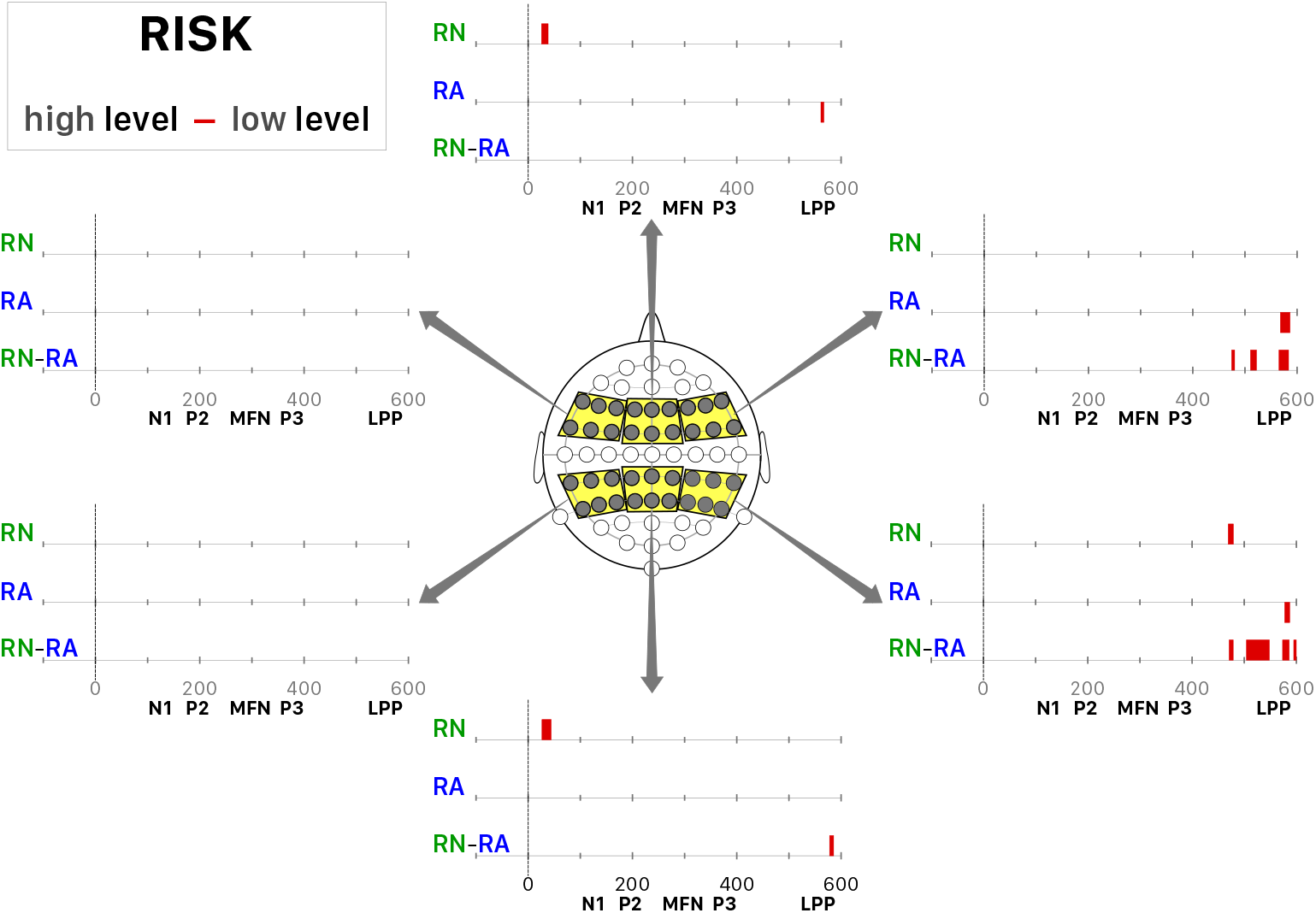
Time course and spatial distribution of the effect of factor “Risk”. Significant intervals of the differential curves corresponding to *Risk_high_* minus *Risk_low_* conditions for groups RN, RA and the difference RN minus RA. The trials were pooled over levels of expected value and the option (safe or risky) selected by the participants. The red labels at the significant intervals point upward for positive and downward for negative values of the amplitude of the differential curve.

#### 3.3.2 Effect of Selected Choice

Next, we considered the time course and spatial distribution of the effect of factor Decision across all six electrode sets (Figure 8). In the medial and lateral frontal sets, we observed a positive effect of choosing the risky option on the amplitude of the differential curve for group RN during P200 and MFN waves. Group RN also showed an even earlier (delay of about 100 ms) positive effect of the risky choice at the left frontal set. Moreover, the enhancement of P200 by risky decisions was more pronounced for RN than RA in medial and left frontal areas. Thus, selecting a risky option is associated with enhanced frontal activity relatively early in a trial, particularly in risk neutral participants. These findings are in line with the hypothesis that choice consistency (which we found to be elevated in RN) involves ERP components associated with automatized stimulus processing.

**Figure 8:**
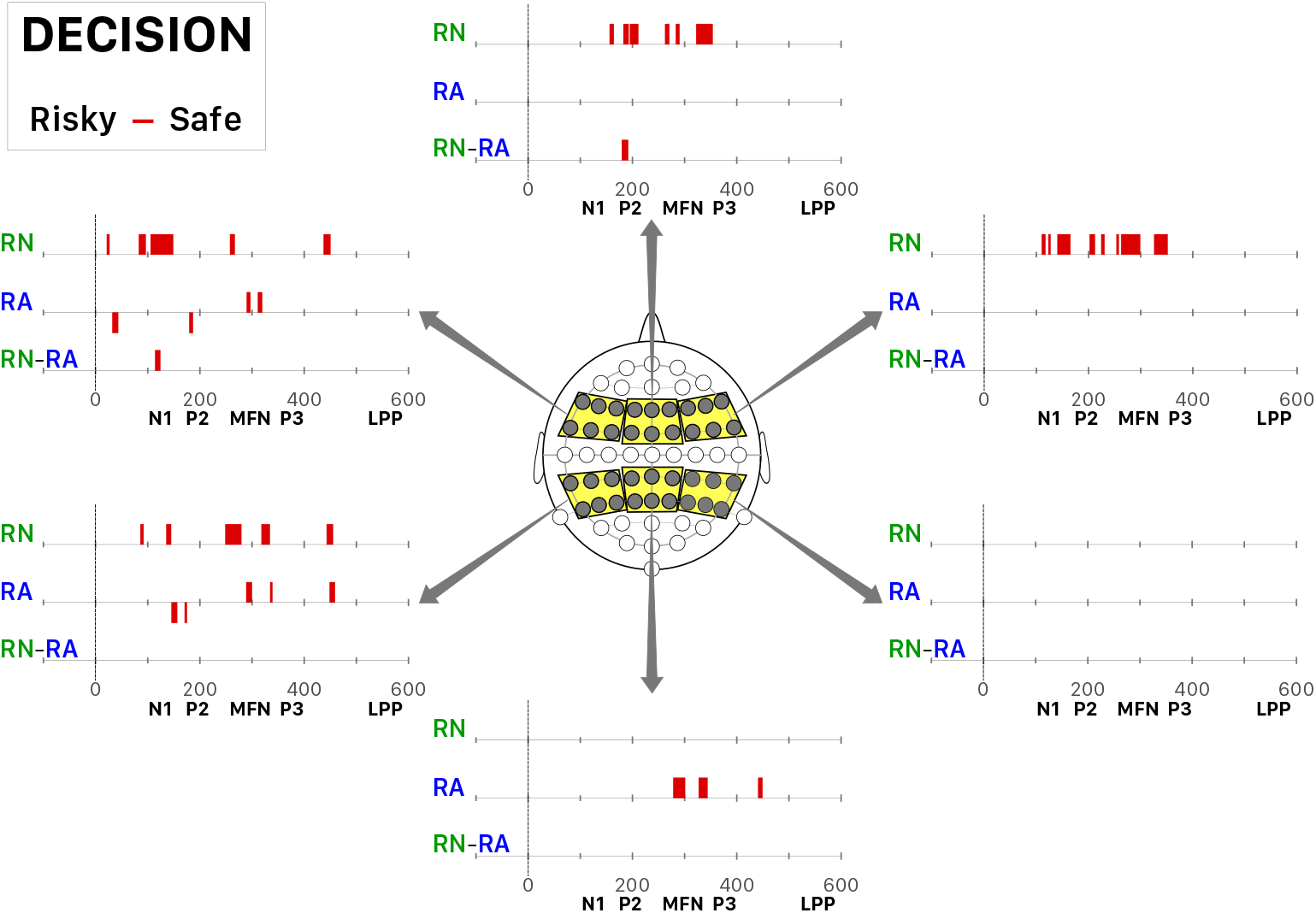
Time course and spatial distribution of the effect of factor “Decision”. Significant intervals of the differential curves corresponding to *Risky* minus *Safe* choices made by the participants for groups RN, RA and the difference RN minus RA. The trials were pooled over levels of expected value and risk. Same legend as Figure 7.

#### 3.3.3 Effect of Magnitude of Expected Value

Along the same line of the previous factors, we analyzed the time course and spatial distribution of factor Expected Value, irrespective of choice taken and risk level (Figure 9). For group RA, we observed only a significant interval at the left frontal electrode set around the time of MFN. In particular, small expected value (EV=30) evoked a stronger negativity than large expected value (EV=60). For group RN, the magnitude of the expected value affected the N1 and P200, but the MFN was not affected in a significant way. We found no significant group differences in the processing of expected value, although the RN group showed more significant effects of expected value overall than the RA group.

**Figure 9:**
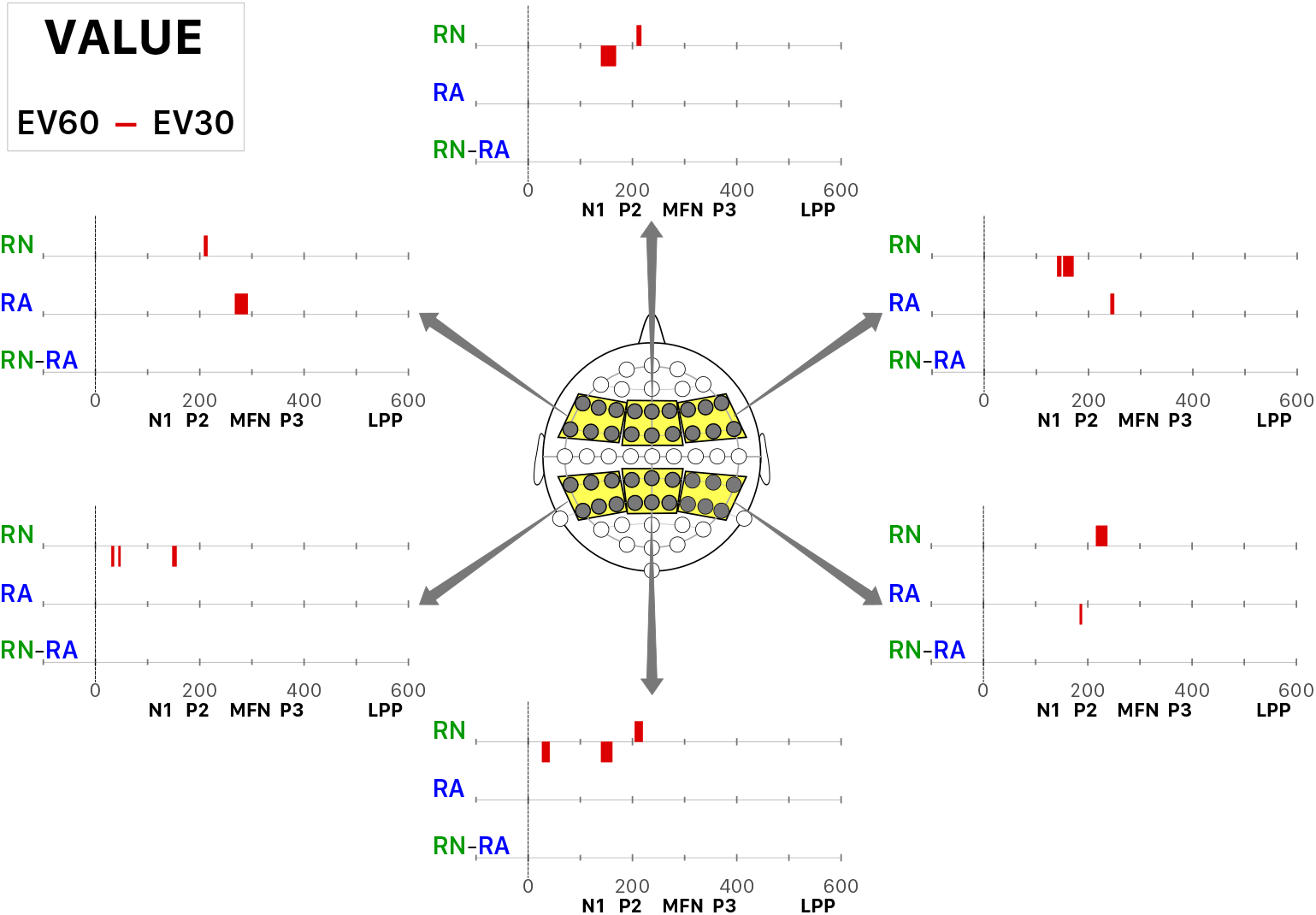
Time course and spatial distribution of the effect of factor “Expected Value”. Significant intervals of the differential curves corresponding to large (EV=60) minus small (EV=30) expected values for groups RN, RA and the difference RN minus RA. The trials were pooled over chosen option (risky or safe) and levels of risk. Same legend as Figure 7.

## 4 Discussion

We investigated the time-resolved neural signatures of risky decision making using the ERP activity elicited by the presentation of choice options. Risk neutral participants showed a greater consistency across trials as compared to risk averse or risk prone participants. Irrespective of risk attitude, about 350 ms after stimulus onset, right parietal electrodes distinguished risky from safe choices. Moreover, both parietal and frontal regions showed differential choice- and risk-related activity early (N1, P200) and late (LPP) during the trial, depending on whether participants were risk neutral or risk averse. A few studies have suggested that stronger cognitive control (i.e., reduced automatic processing) is associated with reduced parietal N1 and frontal P200 (Kehrer et al., 2009; Correa et al., 2020). Along these lines, we may infer that consistency (which was reduced in both risk-averse and risk-seeking participants) is the result of an automatized process.

Expected value appeared to be processed at medium latency components for the RA participants, while it was encoded by the early components in the RN participants. Conversely, risk processing in the positive late potential (LPP) on the right hemisphere differed between RA and RN participants. These findings converge with, and extend earlier reports that expected value is coded earlier than risk (single unit recordings: (Fiorillo et al., 2003); fMRI: (Preuschoff et al., 2006)). Moreover, it is noteworthy that RA participants show value and decision coding in the same area (i.e., the left frontal areas) at the same MFN latency. Based on this finding, it can be hypothesized that stronger EV coding at this latency in left frontal areas helps RA participants overcome their risk aversion.

### 4.1 Risk neutrality and choice consistency

We observed that with stronger deviation from risk neutrality (*RA*_mean_≈0), consistency in the behavior of participants decreased (for both RA and RS groups). We also observed an increase in the mean amplitude of early ERP components for *Risk-neutral* participants compared to *Risk-averse* participants. Two different non-mutually exclusive interpretations can be drawn from this result. First, we confirm that the N1 amplitude at parietal electrodes was influenced by attention factors (Eimer, 1998; Hillyard et al., 1998). This result agrees with the notion that the level of attention paid to stimuli was higher for highly consistent (i.e., RN) individuals. Hence, inconsistency in behavioral performance could be a consequence of inattention. Second, the frontal P200 could reflect the adoption of automated stimulus assessment processes (Luck and Hillyard, 1994; Testa et al., 2020), which could make behavioral choices more consistent across trials. By extension, both risk aversion and risk proneness might characterize individuals whose approach to decision making is based on relatively fuzzy heuristics and/or relatively inaccurate representations of the value of different options. In contrast, the adoption of more automatized heuristics and more accurate value representations might lead to risk-neutral behavior.

In RN participants, we observed that expected value coding was associated with a significantly smaller differential curve in the frontal P200. This differential activity pattern was strongest at the midline for both frontal and parietal electrode sets, which is consistent with differential activations in regions associated with value representations, such as ACC and the prefrontal cortex (PFC) (Noonan et al., 2011; Levy, 2017; Voigt et al., 2020). This result supports the hypothesis that frontal P200 may index expected value representations in the RN group of participants. More generally, our results are in line with the notion that choice inconsistency is (at least in part) related to the variability of the assessment of value (Webb et al., 2019; Kurtz-David et al., 2019).

### 4.2 Late value modulation and differences in risk attitude

Our results support the hypothesis that differences in late risk processing are associated with differences in the attitude towards risk. In the right hemisphere, the impact of risk level was clearly different for RA and RN participants at latencies corresponding to the LPP component. Specifically, RN participants showed a greater mean amplitude for high-risk than low-risk options at parietal electrodes, while RA participants showed the opposite pattern at frontal and parietal electrodes. These results extend those of Wang et al. (2019), who found a larger LPP for options with high probability of realization (i.e., low-risk options) compared to options with low probability (i.e., high-risk options). During decision making requiring self-control, a comparable late effect has been proposed to reflect a top-down modulation of value signals computed in the ventromedial prefrontal region by the dorsolateral prefrontal cortex (Harris et al., 2013). Accordingly, there may be a correspondence between LPP effects found in ERP studies and the lPFC activity found in fMRI and fNIRS studies (Tobler et al., 2007, 2009; Holper et al., 2014). in that risk modulates subjective value signals in both modalities in a risk attitude-specific way. Moreover, a positive correlation between lPFC activity to safer options and the degree of risk aversion may reflect an inhibitory role for lPFC for decision making (Christopoulos et al., 2009). Our findings suggest that such an inhibitory role may be expressed particularly in the LPP activity induced by safer options in RA participants. Further research should use imaging approaches combining high temporal resolution with high spatial resolution to precisely pinpoint the neural activities that underlie the modulation of risk-based value signals in different risk attitude profiles.

### 4.3 Decision-dependent signals

We observed that the difference between the amplitudes of ERPs triggered by risky and safe choices was negative in RA participants at early latencies (<200 ms) in the left hemisphere of RA participants. These results are consistent with the hypothesis (Harris et al., 2013) that rapid stimulus evaluation, as indexed by the P200, plays a key role for decision making and accounts for the difference in behavior between the two groups. Moreover, when all participants were pooled together, we observed that the differential curve was positive for P300 at the right parietal area. This effect also appeared at comparable latencies at medial parietal electrodes for participants in the RA group and at the left parietal set for RA and RN participants. These findings are in agreement with the literature on risky decision making which has shown an enhancement of the P300 amplitude in selecting risky options (Chandrakumar et al., 2018).

### 4.4 Limitations

The biggest limitation of the current study is the low number of participants for an individual difference study. An *a priori* power analysis was conducted to compute the required sample size. This analysis indicated we need a sample size of 23 to achieve a power of 0.95 (1 – *β* error probability) for two-way ANOVA analyses, such as those in Table 2, with one two-level factor (e.g., factor Decision) and one three-levels factor (i.e. Position) and using a desired effect size of 0.9 (to represent a large effect). Moreover, similar to previous studies (Tobler et al., 2007, 2009; Holper et al., 2014), the number of risk-seeking participants was about half that of RA and RN groups. Accordingly, we were not able to further pursue the interesting non-linear trend between risk attitude and choice consistency and relate risk-proneness to meaningful neural signatures. In future work addressing risk-seeking behavior, the total number of participants should be nearly 120 so as to allow for the collection of enough data from the expected number of RS participants.

Another limitation of this study arises from the fact that the SO amounts proposed at risk-free options were distributed according to a pseudo-normal distribution centered on the average certainty equivalents reported by Holper and colleagues (Holper et al., 2017). This manipulation ensures a similar number of risky and safe choices on average and presentation of the same SO for all participants. However, by design, it imposes that risk averse individuals choose the safe options more often than risk tolerant individuals. The comparison between risky and safe choices may suffer from this imbalance. One possibility to address the issue is to tailor the safe options to each individual’s certainty equivalent (Christopoulos et al., 2009), but this has the disadvantage of using different safe options for different participants, which makes it hard to compare option-induced responses. A further limitation of our study is the limited spatial resolution of EEG. It is therefore difficult to relate our results to fMRI reports that pinpoint particular regions with a role in processing individual differences in risk perception or valuation.

## 5 Conclusions

Risk attitudes during decision making vary greatly between individuals. In our task, relative risk neutrality was associated with high consistency across trials, which may reflect a different approach to risky decision making than that used by participants who devalue risk. Neural signatures of the difference between RA and RN groups were observed both at early latencies, thought to reflect differences in attention and stimulus representation, and at longer latencies, thought to reflect more complex valuation processes, decision-making, and/or affective responses. Overall, this study identifies time-resolved components relevant for risky decisions and suggests that examination of individual differences in attitudes toward risk and/or choice consistency is essential in the study of risky decision making, as these individual differences are associated with different brain dynamics.

## 7 Author contributions statement

M.E.J., A.E.P.V. and P.N.T. conceived the experiment(s), M.E.J. and A.L. conducted the experiment(s), M.E.J., G.G., M.F., K.A.L., P.N.T. and A.E.P.V. analyzed the results. M.E.J., A.L., G.G., M.F., K.A.L., P.N.T. and A.E.P.V. wrote and reviewed the manuscript.

## 8 Acknowledgments

We acknowledge financial support by the Swiss National Science Foundation (SNSF), grant POLAP1_178329 to M.E.J. P.N.T was supported by the SNSF (grants 10001C_188878, 100019_176016, and 100014_165884). G.G., M.F., and K.A.L. were supported by grant RF1 AG062666 from the National Institute on Aging.

## References

Bargh, J. A. and Chartrand, T. L. (1999). The unbearable automaticity of being. Am Psychol, 54(7):462–479.

Blavatskyy, P. R. and Pogrebna, G. (2010). Models of stochastic choice and decision theories: Why both are important for analyzing decisions. J. Appl. Econom., 25(6):963–986.

Bossaerts, P. (2010). Risk and risk prediction error signals in anterior insula. Brain Struct Funct, 214(5-6):645–53.

Botvinick, M. M., Braver, T. S., Barch, D. M., Carter, C. S., and Cohen, J. D. (2001). Conflict monitoring and cognitive control. Psychol Rev, 108(3):624–652.

Buysse, D. J., Reynolds, 3rd, C. F., Monk, T. H., Berman, S. R., and Kupfer, D. J. (1989). The Pittsburgh Sleep Quality Index: a new instrument for psychiatric practice and research. Psychiatry Res, 28(2):193–213.

Chandrakumar, D., Feuerriegel, D., Bode, S., Grech, M., and Keage, H. A. D. (2018). Event-Related Potentials in Relation to Risk-Taking: A Systematic Review. Front Behav Neurosci, 12:111.

Christopoulos, G. I., Tobler, P. N., Bossaerts, P., Dolan, R. J., and Schultz, W. (2009). Neural correlates of value, risk, and risk aversion contributing to decision making under risk. J Neurosci, 29(40):12574–83.

Coles, M. G., Gratton, G., Bashore, T. R., Eriksen, C. W., and Donchin, E. (1985). A psychophysiological investigation of the continuous flow model of human information processing. J Exp Psychol Hum Percept Perform, 11(5):529–53.

Correa, Á., Alguacil, S., Ciria, L. F., Jiménez, A., and Ruz, M. (2020). Circadian rhythms and decision-making: a review and new evidence from electroencephalography. Chronobiol Int, 37(4):520–541.

Donchin, E. and Coles, M. G. H. (1988). Is the P300 component a manifestation of context updating? Behav Brain Sci, 11:357–427.

Eimer, M. (1998). Mechanisms of Visuospatial Attention: Evidence from Event-related Brain Potentials. Vis Cogn, 5(1-2):257–286.

Emmorey, K., Midgley, K. J., Kohen, C. B., Sehyr, Z. S., and Holcomb, P. J. (2017). The N170 ERP component differs in laterality, distribution, and association with continuous reading measures for deaf and hearing readers. Neuropsychologia, 106:298–309.

Fecteau, S., Knoch, D., Fregni, F., Sultani, N., Boggio, P., and Pascual-Leone, A. (2007). Diminishing risk-taking behavior by modulating activity in the prefrontal cortex: a direct current stimulation study. J Neurosci, 27(46):12500–5.

Fernandes, C., Pasion, R., Gonçalves, A. R., Ferreira-Santos, F., Barbosa, F., Martins, I. P., and Marques-Teixeira, J. (2018). Age differences in neural correlates of feedback processing after economic decisions under risk. Neurobiol Aging, 65:51–59.

Fiorillo, C. D., Tobler, P. N., and Schultz, W. (2003). Discrete coding of reward probability and uncertainty by dopamine neurons. Science, 299(5614):1898–902.

Frey, D., Johnson, E. D., and De Neys, W. (2018). Individual differences in conflict detection during reasoning. Q J Exp Psychol (Hove), 71(5):1188–1208.

Gehring, W. J. and Willoughby, A. R. (2002). The medial frontal cortex and the rapid processing of monetary gains and losses. Science, 295(5563):2279–2282.

Gratton, G., Coles, M. G., and Donchin, E. (1983). A new method for off-line removal of ocular artifact. Electroencephalogr Clin Neurophysiol, 55(4):468–484.

Grether, D. M. and Plott, C. R. (1979). Economic Theory of Choice and the Preference Reversal Phenomenon. Am. Econ. Rev., 69(4):623–638.

Grothendieck, G. (2013). nls2: Non-linear regression with brute force. R package version 0.2.

Gu, R., Zhang, D., Luo, Y., Wang, H., and Broster, L. S. (2018). Predicting risk decisions in a modified Balloon Analogue Risk Task: Conventional and single-trial ERP analyses. Cogn Affect Behav Neurosci, 18(1):99–116.

Hajcak, G., Dunning, J. P., and Foti, D. (2009). Motivated and controlled attention to emotion: time-course of the late positive potential. Clin Neurophysiol, 120(3):505–10.

Harless, D. W. and Camerer, C. F. (1994). The Predictive Utility of Generalized Expected Utility Theories. Econometrica, 62(6):1251–1289.

Harris, A., Hare, T., and Rangel, A. (2013). Temporally dissociable mechanisms of self-control: early attentional filtering versus late value modulation. J Neurosci, 33(48):18917–31.

Hey, J. D. (2001). Does repetition improve consistency? Exp Econ, 4(1):5–54.

Hey, J. D. and Orme, C. (1994). Investigating generalizations of expected utility theory using experimental data. Econometrica, 62(6):1291–1326.

Hillyard, S. A., Teder-Sälejärvi, W. A., and Münte, T. F. (1998). Temporal dynamics of early perceptual processing. Curr Opin Neurobiol, 8(2):202–10.

Holm, S. (1979). A simple sequentially rejective multiple test procedure. Scand J Statist, pages 65–70.

Holper, L., Van Brussel, L. D., Schmidt, L., Schulthess, S., Burke, C. J., Louie, K., Seifritz, E., and Tobler, P. N. (2017). Adaptive Value Normalization in the Prefrontal Cortex Is Reduced by Memory Load. eNeuro, 4(2):0365.

Holper, L., Wolf, M., and Tobler, P. N. (2014). Comparison of functional near-infrared spectroscopy and electrodermal activity in assessing objective versus subjective risk during risky financial decisions. Neuroimage, 84:833–42.

Intriligator, M. D. (1973). A probabilistic model of social choice. Rev Econ Stud, 40(4):553–560.

Kachelmeier, S. J. and Shehata, M. (1992). Examining Risk Preferences Under High Monetary Incentives: Experimental Evidence from the People’s Republic of China. Am Econ Rev, 82(5):1120–1141.

Kahneman, D. and Tversky, A. (1979). Prospect theory: An analysis of decision under risk. Econometrica, 47(2):263–292.

Kehrer, S., Kraft, A., Irlbacher, K., Koch, S. P., Hagendorf, H., Kathmann, N., and Brandt, S. A. (2009). Electrophysiological evidence for cognitive control during conflict processing in visual spatial attention. Psychol Res, 73(6):751–61.

Knoch, D., Gianotti, L. R. R., Pascual-Leone, A., Treyer, V., Regard, M., Hohmann, M., and Brugger, P. (2006). Disruption of right prefrontal cortex by low-frequency repetitive transcranial magnetic stimulation induces risk-taking behavior. J Neurosci, 26(24):6469–72.

Kroenke, K., Spitzer, R. L., and Williams, J. B. (2001). The PHQ-9: validity of a brief depression severity measure. J Gen Intern Med, 16(9):606–13.

Kurtz-David, V., Persitz, D., Webb, R., and Levy, D. J. (2019). The neural computation of inconsistent choice behavior. Nat Commun, 10(1):1583.

Levy, I. (2017). Neuroanatomical Substrates for Risk Behavior. Neuroscientist, 23(3):275–286.

Logan, G. D. (1988). Toward an instance theory of automatization. Psychol Rev, 95(4):492–527.

Lopez-Calderon, J. and Luck, S. J. (2014). ERPLAB: an open-source toolbox for the analysis of event-related potentials. Front Hum Neurosci, 8:213.

Louie, K., Glimcher, P. W., and Webb, R. (2015). Adaptive neural coding: from biological to behavioral decision-making. Curr Opin Behav Sci, 5:91–99.

Louie, K., Khaw, M. W., and Glimcher, P. W. (2013). Normalization is a general neural mechanism for context-dependent decision making. Proc Natl Acad Sci U S A, 110(15):6139–44.

Luce, M. F., Payne, J. W., and Bettman, J. R. (1999). Emotional Trade-Off Difficulty and Choice. J. Mark. Res., 36:143–159.

Luck, S. J. and Hillyard, S. A. (1994). Electrophysiological correlates of feature analysis during visual search. Psychophysiology, 31(3):291–308.

Luck, S. J., Stewart, A. X., Simmons, A. M., and Rhemtulla, M. (2021). Standardized measurement error: A universal metric of data quality for averaged event-related potentials. Psychophysiology, 58(6):e13793.

Mair, P. and Wilcox, R. (2020). Robust statistical methods in R using the WRS2 package. Behav Res Methods, 52(2):464–488.

Mattsson, L.-G. and Weibull, J. W. (2002). Probabilistic choice and procedurally bounded rationality. Games Econ Behav, 41(1):61 – 78.

McFadden, D. L. (2005). Revealed stochastic preference: a synthesis. Econ. Theory, 26(2):245–264.

Nieuwenhuis, S., Aston-Jones, G., and Cohen, J. D. (2005). Decision making, the P3, and the locus coeruleus-norepinephrine system. Psychol Bull, 131(4):510–532.

Noonan, M. P., Mars, R. B., and Rushworth, M. F. S. (2011). Distinct roles of three frontal cortical areas in reward-guided behavior. J Neurosci, 31(40):14399–412.

Polezzi, D., Sartori, G., Rumiati, R., Vidotto, G., and Daum, I. (2010). Brain correlates of risky decision-making. Neuroimage, 49(2):1886–1894.

Preuschoff, K., Bossaerts, P., and Quartz, S. R. (2006). Neural differentiation of expected reward and risk in human subcortical structures. Neuron, 51(3):381–90.

Rangel, A., Camerer, C., and Montague, P. R. (2008). A framework for studying the neurobiology of value-based decision making. Nat Rev Neurosci, 9(7):545–56.

Rieskamp, J. (2008). The Probabilistic Nature of Preferential Choice. J Exp Psychol Learn Mem Cogn, 34(6):1446–65.

Ryan, M. (2018). Uncertainty and binary stochastic choice. Econ. Theory, 65(3):629–662.

Schuermann, B., Endrass, T., and Kathmann, N. (2012). Neural correlates of feedback processing in decision-making under risk. Front Hum Neurosci, 6:204.

Seidl, C. (2002). Preference reversal. J. Econ. Surv., 16(5):621–655.

Spitzer, R. L., Kroenke, K., Williams, J. B. W., and Löwe, B. (2006). A brief measure for assessing generalized anxiety disorder: the GAD-7. Arch Intern Med, 166(10):1092–7.

Strasburger, H. (2001). Converting between measures of slope of the psychometric function. Percept Psychophys, 63(8):1348–55.

Testa, G., Buongiorno, F., Rusconi, M. L., Mapelli, D., Vettor, R., Angeli, P., Amodio, P., and Schiff, S. (2020). ERP correlates of cognitive control and food-related processing in normal weight and severely obese candidates for bariatric surgery: Data gathered using a newly designed Simon task. Biol Psychol, 149:107804.

Tobler, P. N., Christopoulos, G. I., O’Doherty, J. P., Dolan, R. J., and Schultz, W. (2009). Risk-dependent reward value signal in human prefrontal cortex. Proc Natl Acad Sci U S A, 106(17):7185–90.

Tobler, P. N., O’Doherty, J. P., Dolan, R. J., and Schultz, W. (2007). Reward value coding distinct from risk attitude-related uncertainty coding in human reward systems. J Neurophysiol, 97(2):1621–32.

Tsetsos, K., Usher, M., and Chater, N. (2010). Preference Reversal in Multiattribute Choice. Psychol Rev, 117(4):1275–93.

Voigt, K., Murawski, C., Speer, S., and Bode, S. (2020). Effective brain connectivity at rest is associated with choice-induced preference formation. Hum Brain Mapp, 41(11):3077–3088.

Wang, G., Li, J., Wang, P., Zhu, C., Pan, J., and Li, S. (2019). Neural dynamics of processing probability weight and monetary magnitude in the evaluation of a risky reward. Front Psychol, 10:554.

Webb, R., Levy, I., Lazzaro, S. C., Rutledge, R. B., and Glimcher, P. W. (2019). Neural random utility: Relating cardinal neural observables to stochastic choice behavior. J. Neurosci. Psychol. Econ., 12(1):45–72.

World Medical Association. (2001). World Medical Association Declaration of Helsinki. Ethical principles for medical research involving human subjects. Bull World Health Organ, 79(4):373–374.

Xu, S., Pan, Y., Wang, Y., Spaeth, A. M., Qu, Z., and Rao, H. (2016). Real and hypothetical monetary rewards modulate risk taking in the brain. Sci Rep, 6:29520.

Zhang, D., Gu, R., Broster, L. S., Jiang, Y., Luo, W., Zhang, J., and Luo, Y.-J. (2014). Linking brain electrical signals elicited by current outcomes with future risk decision-making. Front Behav Neurosci, 8:84.

Zheng, Y., An, T., Li, Q., and Xu, J. (2020). Distinct electrophysiological correlates between expected reward and risk processing. Psychophysiology, 57(10):e13638.

Zheng, Y., Li, Q., Wang, K., Wu, H., and Liu, X. (2015). Contextual valence modulates the neural dynamics of risk processing. Psychophysiology, 52(7):895–904.

